# Do adolescents take more risks? Not when facing a novel uncertain situation

**DOI:** 10.1101/206128

**Authors:** Éltető Noémi, Janacsek Karolina, Kóbor Andrea, Takács Ádám, Tóth-Fáber Eszter, Németh Dezso

## Abstract

In real-life decision-making, sub-optimal risk-taking seems characteristic of adolescents. Such behavior increases the chance of serious negative, and at times, irreversible outcomes for this population (e.g., road traffic accidents, addictions). We are still lacking conclusive evidence, however, for an inverted U-shaped developmental trajectory for risk-taking. This raises the question whether adolescents are really more risk-prone or when facing a novel risky situation, they behave just as children and adults do. To answer this question, we used the Balloon Analogue Risk Task (BART) to assess the risky decision making of 188 individuals ranging in age from 7 to 30. The BART provided useful data for characterizing multiple aspects of risk-taking. Surprisingly, we found that adolescents were not more inclined to take risks than children or young adults. Participants in all age groups were able to adapt their learning processes to the probabilistic environment and improve their performance during the sequential risky choice. There were no age-related differences in risk-taking at any stage of the task. Likewise, neither negative feedback reactivity nor overall task performance distinguished adolescents from the younger and older age groups. Our findings prompt 1) methodological considerations about the validity of the BART and 2) theoretical debate whether the amount of experience on its own may account for age-related changes in real-life risk-taking, since risk-taking in a novel and uncertain situation was invariant across developmental stages.

## 1 Introduction

Adolescence is often marked by risky behavior (eg. Eisner, 2002; Sweeten, Piquero, & Steinberg, 2013). While reports about everyday behaviors (such as accident and crime statistics) continuously demonstrate and prove the adolescent peak in risk-taking, data from representative samples in controlled laboratory conditions have started to accumulate only recently. Typically, the occurrence of risk-taking follows an inverted U-shape pattern across development, being relatively low in childhood, increasing and peaking in adolescence, and declining again thereafter (for overviews, see, e.g., Reyna and Farley, 2006; Steinberg, 2004). This developmental pattern is explained by the ‘dual systems model’ (Somerville et al., 2010; Steinberg, 2010), which posits that risk-taking is determined by the interaction of brain systems underlying reward seeking and self-regulation across multiple developmental stages (Galvan, Hare, Voss, Glover, & Casey, 2007) and cultures (Duell et al., 2016, 2017). Along these lines, higher risk-taking in adolescence is predicted by the ‘maturational imbalance’ (Casey, Jones, & Somerville, 2011) between heightened reward sensitivity and immature impulse control (e.g., Braams, van Duijvenvoorde, Peper, & Crone, 2015; McCormick & Telzer, 2017; Peper, Braams, Blankenstein, Bos, & Crone, 2018). Although the motivation of this theory was based on developmental differences, empirical results are still contradictory (Gladwin et al., 2011; Defoe, Dubas, Figner, & Aken, 2015). Not all experiments succeed to confirm elevated risk-taking among adolescents (e.g., Crone et al., 2008; Weller et al., 2010; Paulsen et al, 2011), or showing lower risk-taking behavior in the adolescent groups even in affect-charged tasks (Steinberg et al., 2008). The role of risk opportunity might explain why adolescents do not necessarily assume more risks (Steinberg, 2014). This consideration led to the elaboration of the neuroecological model of risk-taking (Defoe et al., 2015), which predicts more risk-taking among young adults, depending on the social context (e.g., social norms). Some studies showed that both the unconscious appraisal of risk (Shulman & Cauffman, 2014) and risk-taking behavior (Duell et al., 2017) peak in the early 20s. Others have found that young adults do not necessarily assume more risks as their opportunities expand (e.g. Eisner, 2002; Willoughby et al., 2014). Thus, adolescent risk-taking remains puzzling, and more data is needed to reveal those crucial aspects of risky situations that produce the real-world developmental pattern. Here we aimed to map ontogenetic changes of risk-taking in a wide developmental window and used a paradigm that is equally applicable in different age groups and allows for characterizing various aspects of behavior in uncertain risky situations.

The Balloon Analogue Risk Task (BART) (Lejuez et al., 2002) has been used for the empirical assessment of the developmental changes in risk-taking. The BART is an incentivized risk-taking task with immediate outcome feedback on rewards and losses. In the task, people are required to make repeated choices where risk levels may be escalated or eliminated as a result of one’s previous decisions. Thus, the BART provides a link to decision-making accounts of risky behaviors dealing with sequential outcomes (e.g., gradually increasing alcohol intake or driving speed). Indeed, performance on the BART has been shown to be related to these kind of real-world risk-taking behaviors (Lejuez, Aklin, Zvolensky, & Pedulla, 2003; Aklin, Lejuez, Zvolensky, Kahler, & Gwadz, 2005; for an opposite view see Frey et al., 2017). Among adolescents of 11 to 15 years of age, self-reported pubertal status predicted risk-taking on a modified version of the BART beyond relevant demographic characteristics (Collado-Rodriguez, MacPherson, Kurdziel, Rosenberg, & Lejuez, 2014). Also, risk-taking propensity increased across three annual assessment waves in a sample of early adolescents (MacPherson et al., 2010). However, using a cross-sectional design, Humphrey and Dumontheil (2016) were not able to show significant differences in risk-taking behavior on the BART among 12, 15, and 17-year-olds.

As follows, it remains unclear whether there is a behaviorally measurable developmental change in risk-taking. Here, we contribute to the clarification in this field with data from a sample that includes a continuous range of ages that spans preadolescence through young adulthood. Moreover, in this study, we characterize multiple aspects of risk-taking behavior. Aiming to exploit the rich information provided by BART data, we computed both conventional and more refined measures. The latter ones grasp the adaptivity of the participants, reflecting a change in behavior as a function of previous experience and collected information throughout the task. As such, the results of our study may provide a deeper understanding of the developmental course of risk-taking under uncertainty.

## 2 Method

### 2.1 Participants

Two hundred twenty participants between the ages of 7 and 30 took part in the experiment. We excluded participants because of technical problems (three participants), lack of engagement (two participants), atypical amount or quality of sleep relative to their age group or medication use (nine participants), and showing atypical behavior according to Tukey’s (1977) criterion (more than 1.5 times the interquartile range) relative to their age groups along any of the risk-taking measures (twelve participants). One hundred and eighty-eight participants remained in the final sample. Participants were clustered into six age groups between 7-10 (pre-adolescents), 10-13 (early adolescents), 13-16 (mid-adolescents), 16-18 (late adolescents), 18-23 (young adults), and 23-30 (adults) years of age. Table 1 summarizes the descriptive characteristics of the sample. Participants were recruited from local schools with no special curriculum, and universities. The average years of parental education was 14.65±.23 years in the total sample (14.15±.52, 14.57±.71, 15.40±.63, 13.96±.42, 14.45±.71, 15.88±.44, for the six age groups respectively). None of the participants suffered from any developmental, psychiatric, or neurological disorders (based on the parental reports for children and adolescents and based on self-reports for young adults). All participants gave signed informed consent (parental consent was obtained for children), and they received no financial compensation for participation. All experimental procedures were approved by the Institutional Review Board of the university (approval number: 201410; title of the project: Implicit learning and risk-taking across development).

**Table 1.** Descriptive data of demographic and cognitive measures in the whole sample and the five age groups separately.

|  | whole sample | age groups |  |  |  |  |  | F (df) | p |
| --- | --- | --- | --- | --- | --- | --- | --- | --- | --- |
|  | (n = 188) | 7-10 (n = 29) | 10-13 (n = 42) | 13-16 (n = 29) | 16-18 (n = 38) | 18-23 (n = 27) | 23-30 (n = 23) |  |  |
| Gender (male/female) | 80/108 | 14/15 | 21/21 | 15/14 | 16/22 | 8/19 | 6/17 |  |  |
| Age M (SD) | 15.55 (5.14) | 8.91 (0.72) | 11.20 (0.87) | 14.56 (0.83) | 16.91 (0.54) | 20.75 (1.17) | 24.65 (1.21) |  |  |
| Digit span M (SD) | 5.55 (1.13) | 4.72 (0.52) | 5.17 (0.86) | 5.74 (1.12) | 5.68 (1.16) | 6.22 (1.18) | 6.04 (1.22) | 8.44 (5, 181) | < .001 |
| Counting span M (SD) | 3.44 (.86) | 2.68 (0.50) | 3.28 (0.93) | 3.75 (0.86) | 3.46 (0.65) | 3.96 (0.78) | 3.63 (0.80) | 9.46 (5, 180) | < .001 |
| Corsi blocks span M (SD) | 5.12 (1.04) | 4.38 (1.04) | 4.75 (0.70) | 5.58 (1.05) | 5.50 (1.13) | 5.33 (0.96) | 5.22 (0.79) | 7.38 (5, 180) | < .001 |
| Phonemic fluency* M (SD) | 14.33 (5.64) | 8.37 (2.73) | 11.82 (3.12) | 13.75 (4.41) | 15.04 (4.34) | 18.19 (4.61) | 20.58 (6.36) | 27.54 (5, 175) | < .001 |

Note: \* number of correct words

### 2.2 Stimuli, design, and procedure

We used a modified version of the Balloon Analogue Risk Task (BART) (Fein & Chang, 2008), a well-validated and widely-used behavioral measure of risk-taking, originally developed by Lejuez et al. (2002). The structure and appearance of the BART were similar as described in previous studies (Kardos et al., 2016; Kóbor et al., 2015; Takács et al., 2015). The task was implemented in E-Prime 2.0 (Psychology Software Tools, Inc.). This version of the task is more suitable for the current study because of (1) the higher appealing characteristics (incrementing potential reward) that makes the tool more sensitive to individual differences and (2) the shorter trial length that makes the task more applicable for all age groups.

During this task, participants repeatedly decided whether to continue or discontinue inflating a virtual balloon that could grow larger or explode. After each successful pump, the scores in a virtual temporary bank (the accumulated score on a given balloon) increased, as well as the size of the balloon. Instead of further pumping the balloon, participants could have finished the actual balloon trial and collected the accumulated score, which was transferred to the virtual permanent bank. Two response keys on a keyboard were selected either to pump the balloon or to finish the trial. There were two possible outcomes as results of a pump: The size of the balloon together with the score inside increased (positive feedback) or the balloon burst (negative feedback). The balloon burst ended the actual trial, and the accumulated score on that balloon was lost, but this negative event did not decrease the score in the permanent bank.

Five information chunks persistently appeared on the screen during the task: (1) the accumulated score for a given balloon in the middle of the balloon, (2) the score in the permanent bank, (3) the score collected from the previous balloon, (4) the response key option for pumping the balloon, and (5) the other response option for collecting the accumulated score. After collecting the accumulated score that ended the balloon trial, a separate screen indicated the gained score. This screen or the other one presenting balloon burst was followed by the presentation of a new empty (small-sized) balloon indicating the beginning of the next trial (Figure 1). Participants had to inflate 30 balloons in this version of the BART.

**Figure 1.**
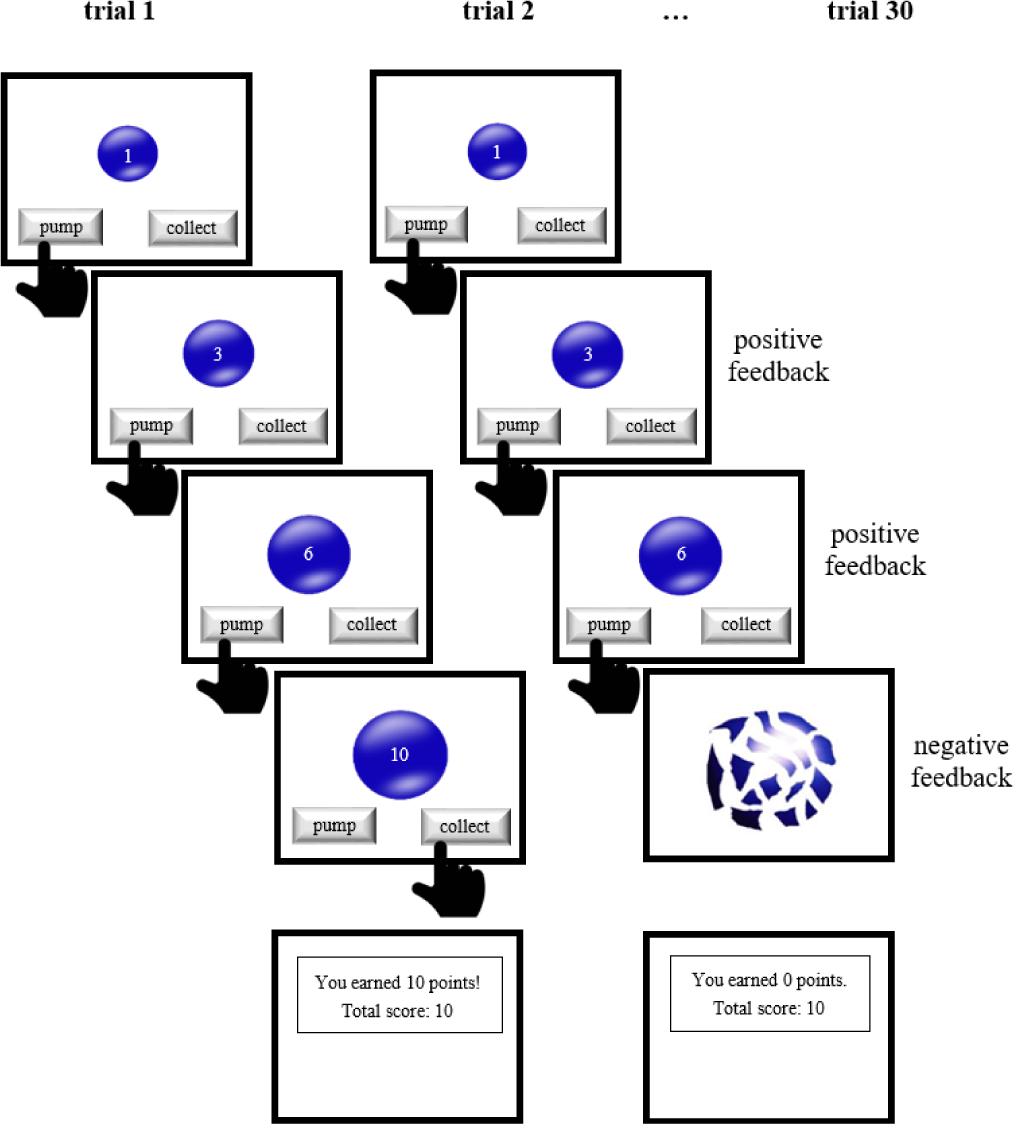
Schematic diagram for the BART. The BART is an ecologically valid model for the assessment of risk-taking behavior. During this task, participants repeatedly decide whether to continue or discontinue inflating a virtual balloon that can grow larger or explode. A larger balloon has both a higher probability of explosion but also a potential for greater reward.

Concerning the structure of the task, each successful pump increased the potential reward but also the probability to lose the accumulated score because of balloon burst. Although the regularity determining balloon bursts was unknown to participants, it followed three principles: (1) balloon bursts for the first and second pumps were disabled; (2) the maximum number of successful pumps for each balloon was 19; (3) the probability of a balloon burst was 1/18 for the third pump, 1/17 for the fourth pump, and so on for each further pump until the 20th, where the probability of a balloon burst was 1/1. Thus, the probability of a balloon burst is given by the formula:

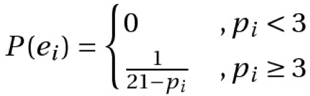

where *P*(*e_i_*) is the probability of the balloon explosion on the *i*^th^ trial (balloon), and *p*_*i*_ is the number of pumps on the *i*^th^ trial (balloon).

One point was added to the temporary bank for the first successful pump, two for the second (i.e., the accumulated score for a given balloon was 3), three for the third (i.e., the accumulated score was 6); thus, the formula for the accumulated trial earning at the point of cashing out is:

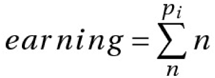

where *p*_*i*_ is the number of pumps on the *i*^th^ trial (balloon), and *n* is the pump index within a trial, (e.g., first, second etc., pump).

According to the instructions, participants were asked to achieve as high score as possible by inflating empty virtual balloons on the screen. Participants did not receive any guaranteed reward. However, they were informed that the participant with the most total earnings in a subgroup of 10 (within the same age group) was rewarded. In order to induce comparable subjective incentive value among age groups, the reward’s identity was unknown to the participants during data acquisition. The reward was a small gift in all age groups and changed according to the age groups.

Besides risk-taking measurement, standard neuropsychological tests were administered. The digit span task (Isaacs & Vargha-Khadem, 1989; Hungarian version: Racsmány, Lukács, Németh, & Pléh, 2005) was used to measure phonological short-term memory capacity. The counting span task (Case, Kurland, & Goldberg, 1982; Hungarian version: Fekete, Filep, Gyüre, Ujvári, Janacsek & Németh, 2010) is a complex working memory task, which requires not only information storage and rehearsal, but also simultaneous processing of additional information. The Corsi blocks task (Kessels, Van Zandvoort, Postma, Kappelle & De Haan, 2000) was used to measure visuospatial short-term memory. The phonemic fluency task (Spreen and Strauss, 1991; for Hungarian version, see Tanczos el al., 2014a, 2014b) is widely used to measure the central executive component of the working memory model (Baddeley, 1992).

The tasks were administered in two sessions in two subsequent days. The BART was administered on the first day, and the neuropsychological tests were administered on the second day. On the second day, the order of the tasks was counterbalanced across participants. Standard instructions and conditions were applied across all age groups. Participants did not receive financial compensation.

### 2.3 Risk-taking measures

The BART allowed us to quantify multiple aspects of risk-taking. We calculated four conventional BART measures: the mean number of pumps, the mean adjusted number of pumps (mean number of pumps on unexploded balloons; see Lejuez et al., 2002), the number of balloon bursts (Schmitz et al., 2016) and the earnings (Koscielniak et al., 2016; Schmitz et al., 2016). In the analyses, we either used these measures averaged across all trials or in three bins (1-10 trials, 11-20 trials, and 21-30 trials), in order to be able to assess changes in risk-taking across the 30 trials (as in Lejuez et al., 2002).

One potential drawback of the conventional measures might be that they do not take into account the actual experience of the participant, which – due to the probabilistic structure of the task – is highly variable even for participants with identical behavior. BART data provides information above and beyond the conventional measures, allowing for the assessment of further important aspects of risk-taking (Schmitz et al., 2016). Therefore, we computed the *post-explosion reactivity* (which in Schmitz et al., 2016 is referred to as Δ post-loss pumps), that is the mean number of pumps relative to the pumps at the previous balloon explosion (or the mean number of pumps in the case of more subsequent bursts), averaged across all explosions in the task, according to the following formula:

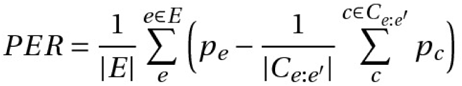

where *E* is the set of trial indices, where the balloon exploded; *e* is an individual trial where the balloon exploded; *C*_*e:e’*_ is a set of trials (balloons) in between two consequent explosions *e* and *e’*; *c* is an individual cash-out trial in *C*_*e:e’*_; *p*_*e*_ denotes the number of pumps on the *e*^th^ explosion trial and *p*_*c*_ denotes the number of pumps on the *c*^th^ cash-out trial.

We also computed *the immediate post-explosion reactivity* measure, where we only took into account the first post-explosion trials:

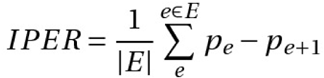

where *p*_*e+1*_ is the number of pumps on the trial (balloon) right after the explosion.

Humphreys & Lee (2011) used an index corresponding to the immediate post-explosion reactivity as a measure of sensitivity to negative punishment. Accordingly, the two measures described above represent the extent to which participants adjust their responses to the negative feedbacks. If the post-explosion reactivity/immediate post-explosion reactivity score is 0, it shows that, on average, the participant inflated to the highest level where the balloon did not explode yet in the latest explosion trial. A negative value indicates less risk-taking, and a positive value indicates more risk-taking than it would be perfectly adapted according to the information from the negative feedback. Note that in the case of these two measures, the bin-wise analysis was not possible, only the conventional measures, since the number of data points taken into account was restricted by the number of balloon bursts.

## 3 Results

First, we assessed whether a continuous quadratic model accounted for the developmental change of BART measures. We found that none of the BART measures – namely, the mean number of pumps, the mean adjusted number of pumps, the number of balloon bursts, the earning, and reactivity measures – followed an inverted U-shape across development (all *p*s >.094, see Table 2 and Figure 2).

**Table 2.** Summary of the results from frequentist regression analysis performed on all BART measures, testing quadratic fits. Note that regression coefficients are not included in the table, since none of the models accounted for a significant amount of variance in risk-taking.

| Measure | Quadratic fit |  |  |
| --- | --- | --- | --- |
| | $R^2$ | $F$ | $p$ |
| Mean Number of Pumps | .008 | .731 | .483 |
| Mean Adjusted Number of Pumps | .002 | .219 | .804 |
| Number of Balloon Bursts | .007 | .650 | .523 |
| Total Earnings | .010 | .926 | .398 |
| Post-Explosion Reactivity | .020 | 1.842 | .161 |
| Immediate Post-Explosion Reactivity | .026 | 2.399 | .094 |

**Figure 2.**
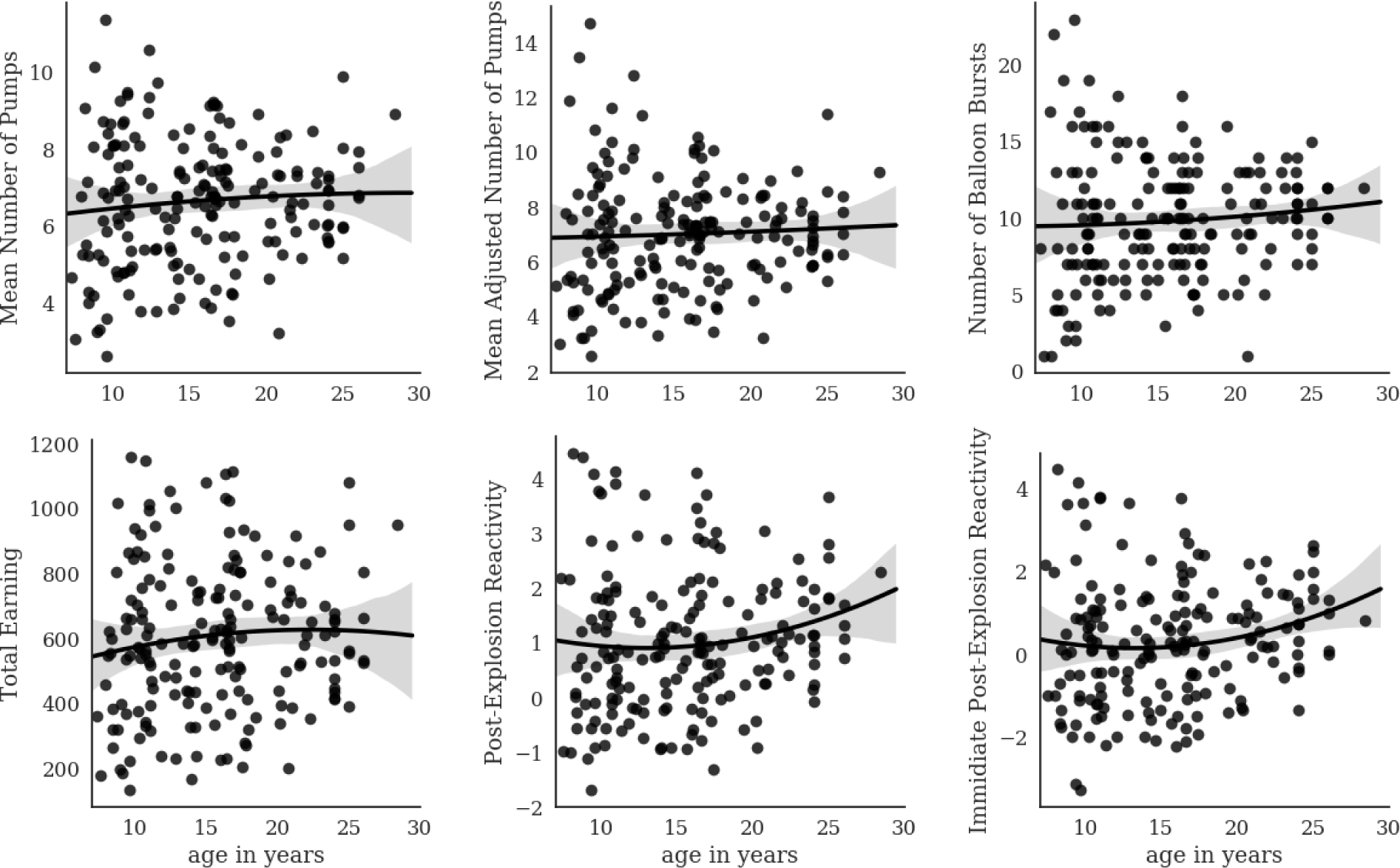
Quadratic regression models fitted to developmental data on six BART measures. None of the models were significant. *Note:* Higher numbers indicate greater risk-taking. Grey bands represent the 95% confidence intervals.

Next, we assessed the developmental modulation of the *pattern* of risk-taking across the 30 trials. For this end, we conducted mixed design ANCOVAs with BIN (1-10 trials, 11-20 trials, 21-30 trials) as the within-subjects factor and AGE GROUP (7-10, 10-13, 13-16, 16-18, 18-23, 23-30) as the between-subjects factor, controlling for verbal, visuospatial, and complex working memory.

The dependent variables were the mean number of pumps, the mean adjusted number of pumps, the number of balloon bursts, and the earnings, respectively. The Greenhouse-Geisser epsilon correction (Greenhouse & Geisser, 1959) was used when necessary. Risk-taking was gradually heightened across the trials as shown by the main effect of BIN (all *p*s < .001) on the mean number of pumps, the mean adjusted a number of pumps, and the earning. The number of balloon bursts, however, did not change across trials (*p* = .328). Most importantly, we found no significant BIN*AGE GROUP interaction (all *p*s > .683, see Table 3 and Figure 3).

**Table 3.**
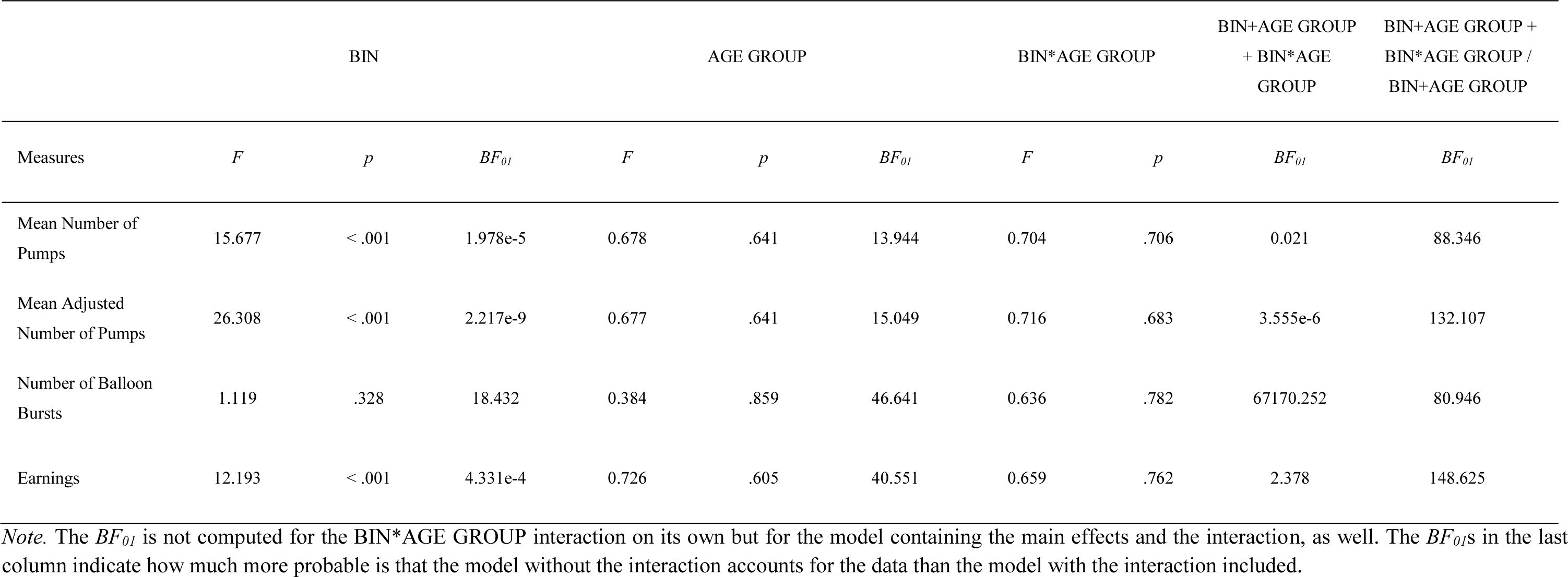
Summary of the results from both frequentist and Bayesian mixed design ANCOVAs performed on all BART measures in three bins of the BART and six age groups, controlling for cognitive measures.

**Figure 3.**
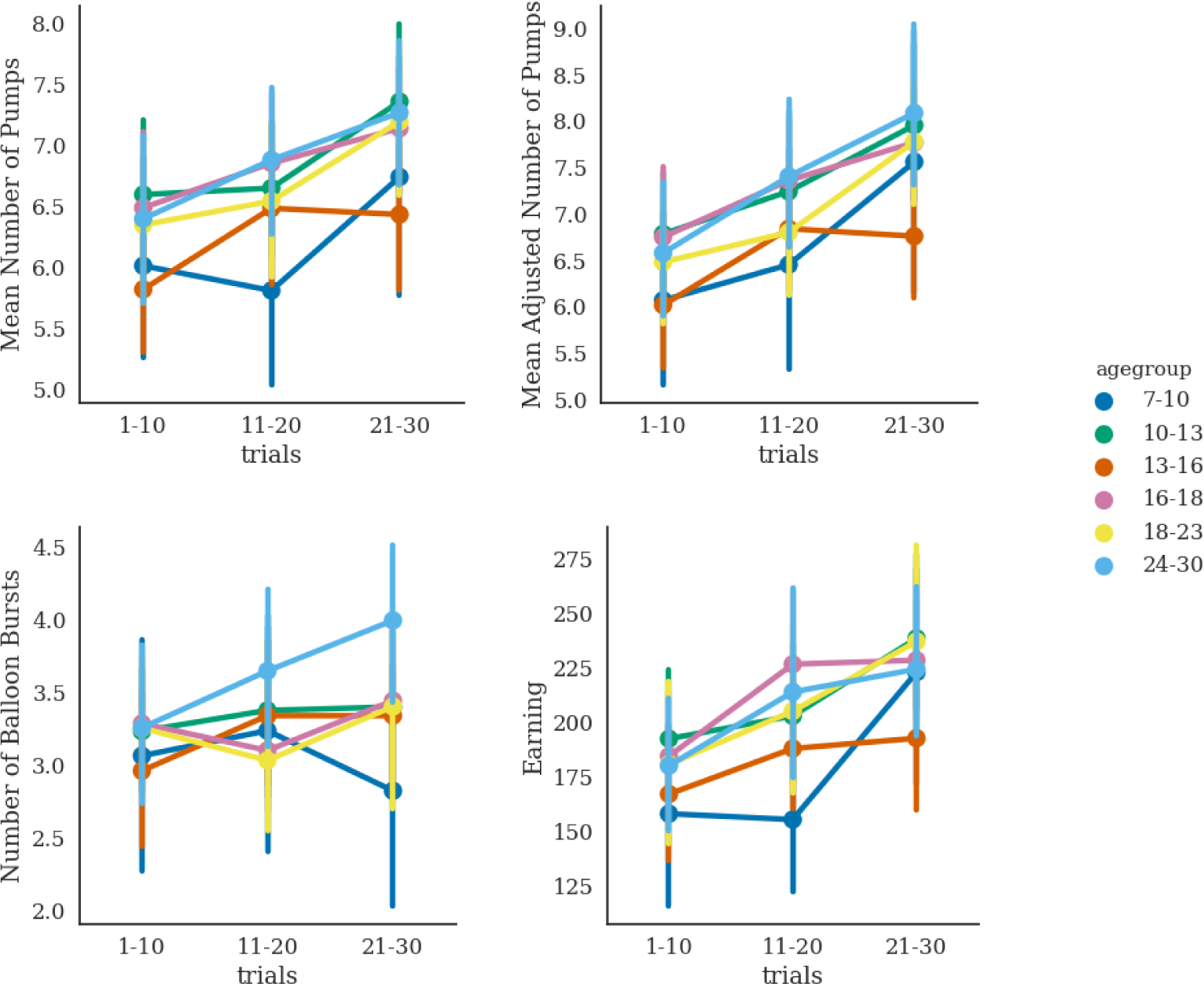
Change in risk-taking across three bins in the BART by age groups (indicated by different colors), as reflected by four measures. The mean number of pumps, the mean adjusted number of pumps, and the earning are higher as the task proceeds, while the number of the balloon bursts remain unchanged. This behavioral pattern characterizes all five age groups, with no significant differences among them. *Note.* Higher numbers indicate greater risk-taking. Error bars represent the 95% confidence intervals.

In order to be able to conclude whether our data supported the lack of developmental changes, we conducted Bayesian repeated measures ANCOVAs, as well. According to Lee and Wagenmakers (2013), values of the Bayes Factor (*BF*_*01*_) larger than 1 indicate anecdotal evidence, BF_01_ values larger than 3 indicate moderate evidence, and BF_01_ values larger than 10 indicate strong evidence for the lack of group differences. We found strong evidence for the lack of age effects on all BART measures (all *BF*_*01*_s > 13.944). The results of the Bayesian mixed design ANCOVAs confirmed the absence of the BIN*AGE GROUP interaction, regarding all BART measures. The models without the interaction effects could account for the data better than the models with the interaction effects (all *BF*_*01*_s for the models containing only the main effects are at least 80.946 times larger than the *BF*_*01*_s for the models containing both the main effects and interaction effects). Moreover, the simpler model containing only the main effect of BIN could account for the data better than the model containing the effects of both BIN and AGE, confirming the lack of age-related changes. Thus, the risk-taking uptrend across trials is characteristic of the sample, irrespectively of developmental phase.

Furthermore, one-way ANCOVAs with AGE GROUP (7-10, 10-13, 13-16, 16-18, 18-23, 23-30) as a between-subject factor on Post-Explosion Reactivity and Immediate Post-Explosion Reactivity, controlling for verbal, visuospatial, and complex working memory. There were no significant differences on any of the BART measures among the age groups (*F*(5, 167) = 1.305, *p* =.264; *F*(5,167) = 1.290, *p* =.270, respectively). Bayesian ANCOVAs supported these results as we found moderate evidence for the lack of group differences (*BF*_*01*_ = 8.553; *BF*_*01*_ = 8.568, respectively).

## 4 Discussion

Despite the common real-life accounts for elevated risk-taking in adolescence, findings on age differences have remained conflicting. Here we aimed to assess risk-taking using the same experimental context that allows for measuring multiple aspects of sequential risk-taking in a relatively long period of ontogeny, from 7 to 30 years of age. We were not able to show age-related changes in risk-taking measures. Moreover, the results of Bayesian analyses proved that the *adevelopmental model* of sequential risk-taking data is the most plausible. Overall, the lack of age-related changes was consistently found for the overall risk-taking as well as for the course of risk-taking across multiple decisions.

Our firmly negative results may be surprising, as they neither fit into the theories of adolescent risk-taking nor fall into the line of empirical results showing an inverted U-shaped developmental course of risk-taking. For example, neurodevelopmental imbalance models suggest that there is a potential for an imbalance between cognitive and affective processes in adolescence (Somerville & Casey, 2010; Steinberg, 2007), and these models postulate, in particular, that in emotionally charged (‘hot’) situations, adolescents’ hypersensitive motivational-affective system often overrides cognitive control capacities that adolescents might have. Indeed, meta-regression analyses revealed that adolescents take more risks than adults on ‘hot’ tasks with immediate outcome feedback on rewards and losses (Defoe et al., 2015). Both MacPherson et al. (2010) and Lejuez et al. (2014) showed that risk-taking elevates across multiple waves of annual assessments in an adolescent sample. Likewise, Peper et al. (2018) found increasing risk-taking across three time-points separated by 2 years, leveling off in young adulthood. However, along with previous studies (Lejuez et al., 2002; Kardos et al., 2016), we showed that risk-taking levels rise across a single session of BART as well. As predicted by the prospect theory (Kahneman & Tversky, 1979), loss aversion leads to sub-optimal performance in the initial phase of the sequence of probabilistic decisions, and most participants approach the optima by accumulating experience. Therefore, it becomes hard to interpret the results of the longitudinal studies mentioned above, since the differences among the annual assessments might reflect both developmental changes and trial-by-trial changes captured within sessions. The flat developmental curve derived from our cross-sectional data prompts to consider the possibility that decision-making processes taking place in uncertain and affect-charged situations might not all be subject to ontogenetic changes. Thinking of human decision-making as Bayesian inference (e.g., Griffiths, Kemp, & Tenenbaum, 2008), it depends on estimated probability distributions based on previous observations, in other words, on the *prior*. Adolescents, having had less experience, have less information available for representing the probabilistic structure of decision situations via *prior*s. Along these lines, we can speculate that differences not only in the decision-making processes per se, but also differences in the reliability of the priors that serve as input to those processes can account for the real-life observations about adolescent risk-taking. By the use of tasks that participants had not had previous experience with, and that are distinct from real-life risk-taking situations, the role of priors is supposedly reduced. In the novel decision context of the BART, age groups did not diverge in terms of risk-taking behavior.

Another possible explanation of our results is of a methodological nature. The lack of developmental changes in our study may suggest that the original version of the BART (Lejuez et al., 2002) is not entirely capable of modeling all crucial components of ‘real-life’ risky situations. Nevertheless, the BART fits to both the *naïve* and *economic* definitions of *risk* in a sense that each successive pump increases the exposure to negative outcome and relates to increased variance in the possible outcomes (Helfinstein et al., 2014), and the exact probability of outcomes (balloon burst or balloon increase) is not known to participants at any moment. Thus, the BART could still be closer to many everyday risk-taking behaviors than other tasks that have been used for investigating the development of risk-taking. These include tasks where the probability of outcomes is either clearly stated to the participants (e.g., Van Duijvenvorde, Jansen, Bredman, & Huezinga, 2012 using the informed version of the Iowa Gambling Task; Van Leijenhorst, Westenberg, & Crone, 2008 and Van Leijenhorst, Moor, de Macks, Rombouts, Westenberg, & Crone, 2010 using a modified version of the Cambridge Gamble Task), or relatively easily discoverable (Huizenga, Crone, & Jansen, 2007). Future studies directly examining and testing how different tasks relate to different aspects of real-life risk-taking seem warranted.

Different versions of the BART might lead to different results as well. In the most recent cross-sectional and cross-national study on the development of risk-taking (Duell et al., 2018), the modified version of the BART was used. They found a curvilinear developmental pattern on the total sample of 5227 participants, the peak in risk-taking occurring in the early to mid-20s. Moreover, the quadratic pattern in risk-taking was significant in 8 out of 11 of the countries studied. In the BART version used by Duell et al., the balloon inflated continuously until the participant paused the inflation by pressing the space bar again. From this point, participants could incrementally inflate the balloon by pressing the space bar. When the desired inflation size was reached, participants hit a separate key to obtain the points accumulated. Thus, on each trial, the participants had to stop the automatic inflation first, and then actively inflate the balloon further. By contrast, in the version that was used both in our study and the original Lejuez et al. (2002) study, the balloons were only inflated as a consequence of the active risk-taking of the participants. That is, there was no automatic ‘startup’ inflation at the beginning of the task. The hybrid BART version used by Duell et al. might be sensitive to other aspects of risk-taking too since each trial involves not only active risk seeking but also active risk aversion in the first step of the trial (i.e., when participants have to stop the automatic inflation). In the present study, using the BART version in which individuals themselves had to assume risks in every step of a trial, as opposed to deciding from which point to start doing so, we did not find the same age-related differences as Duell et al. Notably, the study by Duell et al. was not only methodologically different, but also more extensive, using a sample of size 5227 as opposed to our sample of 188. Nevertheless, power simulations confirmed that the probability of detecting the quadratic pattern reported by Duell et al. on a random sample of size of 188 is above 80% (Figure S3). This leaves us with the explanation that the aforementioned methodological factors contributed to the emergence of the developmental differences in risk-taking in the study by Duell et al. and not ours. Development might differentially influence risk aversion and active risk-taking. Certainly, this remains an open question and it should be addressed with an experiment in which risk aversion and active risk-taking is directly manipulated, by using both BART versions in different age groups.

It is also conceivable that complex strategies and impulsivity influence behavior on the BART. To achieve a larger reward (higher score), participants had to postpone their responses and not to collect smaller but immediate rewards. As such, higher rates of temporal discounting might lead to lower scores. Impulsivity, on the other hand, has been shown to be related to more and faster balloon pumps on the BART (Lejuez et al., 2002), although these results have not been demonstrated consistently (e.g., Hunt, Hopko, Bare, Lejuez, & Robinson, 2005). It is important to consider the possibility that impulsivity has an opposite effect too – an urge to cash out soon. If two behavioral consequences of impulsivity, delay aversion and less deliberative, fast risk-taking cancel each other out in the final decision that is measured in the task, the relationship between risk-taking and impulsivity remains undiscoverable. It is a challenge for future studies to disentangle how impulsivity might contribute to both lower and higher risk-taking than the optimum, due to premature cash-outs and fast over-inflation of the balloons in the BART. Such studies should also use other tasks and questionnaires that specifically measure impulsivity, complex strategies, and risk-taking propensity, in order to clarify the individual and interactive effects of these factors. Despite choosing a BART version that is presumably applicable in developmental studies for its shortness and sensitivity, some design features may have introduced confounding factors. Particularly, the payout scheme (a reward was granted only for the participant with the most total earnings in a subgroup of 10) may have induced complex strategies (e.g., reasoning about other’s probable performance). Whilst in theory, this special payout scheme makes the interpretation of the number of pumps more difficult, based on our and previous studies (e.g., Hunt et al. 2005) we believe that it does not affect the results. On the other hand, the characteristics of the feedback are presumably crucial, since the BART can be viewed as a trial-and-error learning task. Findings suggesting that children’s learning benefits more from positive than negative feedback (Lange-Küttner, Averbeck, Hirsch, Wießner, & Lamba, 2012) should be taken into account when interpreting our and other results on the development of risk-taking on the BART. Further studies should clarify the role of the mere acknowledgment of other’s role in one’s own decision, and the differential effects of positive versus negative feedback in risk-taking across the development.

Peer presence *per se* is proved to have an effect on the decision-making of adolescents, leading to riskier choices (e.g., Chein, Albert, O’Brien, Uckert, & Steinberg, 2011; Smith, Chein, & Steinberg, 2014), or even more cautious behavior, probably depending on the situational context, the reward scheme, and the stochasticity of the task (Kessler, Weichold, Silbereisen, 2017). Aiming to promote the disentanglement of the different factors of risk-taking, here we ruled out the social aspects, allowing for the examination of the cognitive process. Thus, we contribute to the literature with data on how teenagers decide what risks are worth taking when they are not directly influenced by friends or other peers (without implying that the results are decidedly explained by the absence of peer presence). We suggest further refinement of the theories of adolescent risk-taking in order to be able to pinpoint the crucial dispositional and social factors that are developmentally determined.

Taken together, the current findings extend the vast literature on adolescent risk-taking by providing substantial evidence for no developmental changes from 7 to 30 years of age. While most studies in this field employ one or two measures, here we conducted a more thorough assessment of risk-taking by looking separately at the behavior 1) on trials with no negative feedback, 2) the change in behavior after a negative feedback, 3) and overall task performance, in the BART. Despite these and other methodological strengths, our study is not conclusive, thus warranting more extensive data analysis and refinement of theoretical frameworks in this field.

## Supplementary Material

We conducted a power simulation to determine the probability of detecting the quadratic pattern described by Duell et al. (2018) using a sample of size 188. Since the quadratic model has more than one parameter, the analytical solution for power analysis is problematic. Instead, we implemented a computational simulation (Jupyter Notebook available: https://github.com/noemielteto/BART_DEV_power_simulation/). First, we have generated 5227 data points to emulate the empirical total sample collected by Duell et al., based on the parameters of the quadratic function fitted to their sample, reported in Table 4 of their paper. Thus, the formula for generating a BART score was:

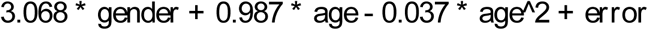

We sampled age uniformly in the range between 0 and 20. This corresponds to the age range between 10 and 30, after centering to 0 (the centering was used in the Duell et al. paper, so we conformed to this in our simulation). The gender term was randomly set to 0 (female) or 1 (male). The error term was adjusted until the standard error of the mean of the unstandardized beta approached the standard error values of the parameters as reported by Duell et al. in Table 4. The resulting sample of 5227 data points reflects the marked quadratic pattern (Figure S1) that was presented in the paper.

Next, we took 10000 random subsamples of N=188 from the total sample of 5227, and we tested the quadratic fit (F-test) of these random subsamples. In Figure S2 we show the first five samples and the p-value of the F-test for the quadratic fit.

The results of this procedure can be interpreted as an analytical solution for power analysis. In Figure S3 we show that the probability of detecting the quadratic pattern reported by Duell et al. on a random subsample of N=188 is above 80%.

Given that most researchers use a threshold for statistical power of 80%, the sample size used in our study meets the consensual standard for adequacy. If the same developmental trajectory of risk-taking had been characteristic to our sample as to the large sample assessed by Duell et al., a sample size of 188 would have allowed us to detect the pattern with a high probability. Yet, the p-values for almost all of our statistical tests presented in the MS are so high that the probability of encountering them given an existing quadratic pattern was virtually 0 in our simulation. In other words, it is improbable that our results are false negatives, as supported by our power simulation.

**Figure S1.**
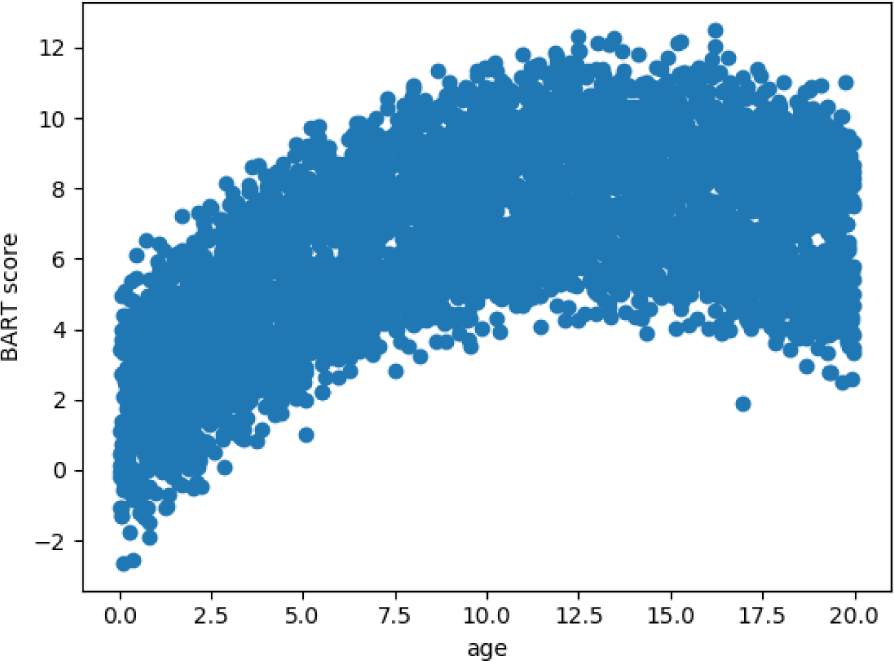
The simulated sample of N = 5227 based on the regression parameters reported by Duell et al. (2018). Note that the X-axis represents the centered age values (ranging from 0 to 20, corresponding to the raw data ranging from 10 to 30).

**Figure S2.**
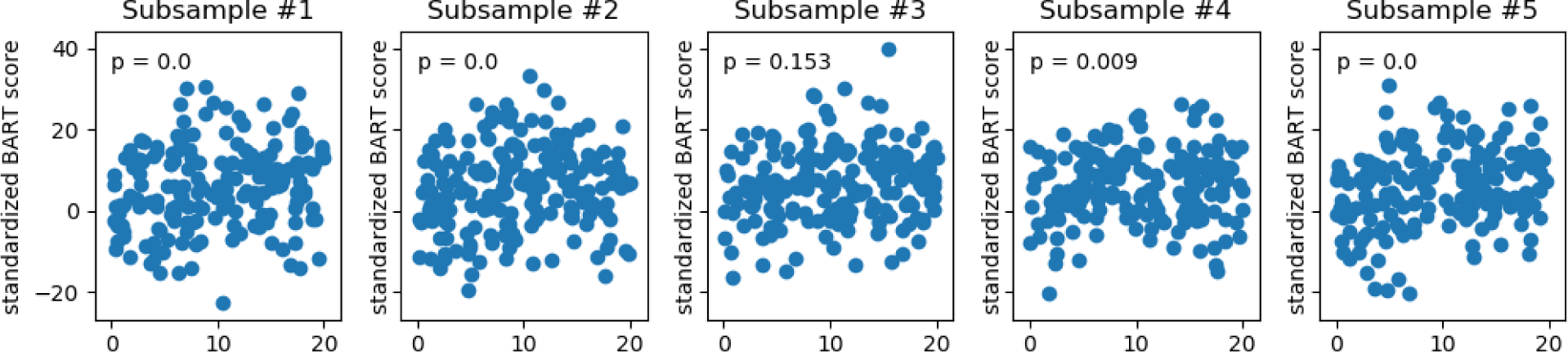
The first five subsamples of N=188 taken from the total sample of N=5227 that represents the developmental effect presented by Duell et al. (2018). The quadratic fit can be detected on nine of the ten subsamples with an alpha value of 0.05, and on eight of the ten samples with an alpha value of 0.01. Note that p = 0.0 indicates p < 0.001. Naturally, different subsamples will be yielded by running the simulation script, but the ratio of significant quadratic fits is stable.

**Figure S3.**
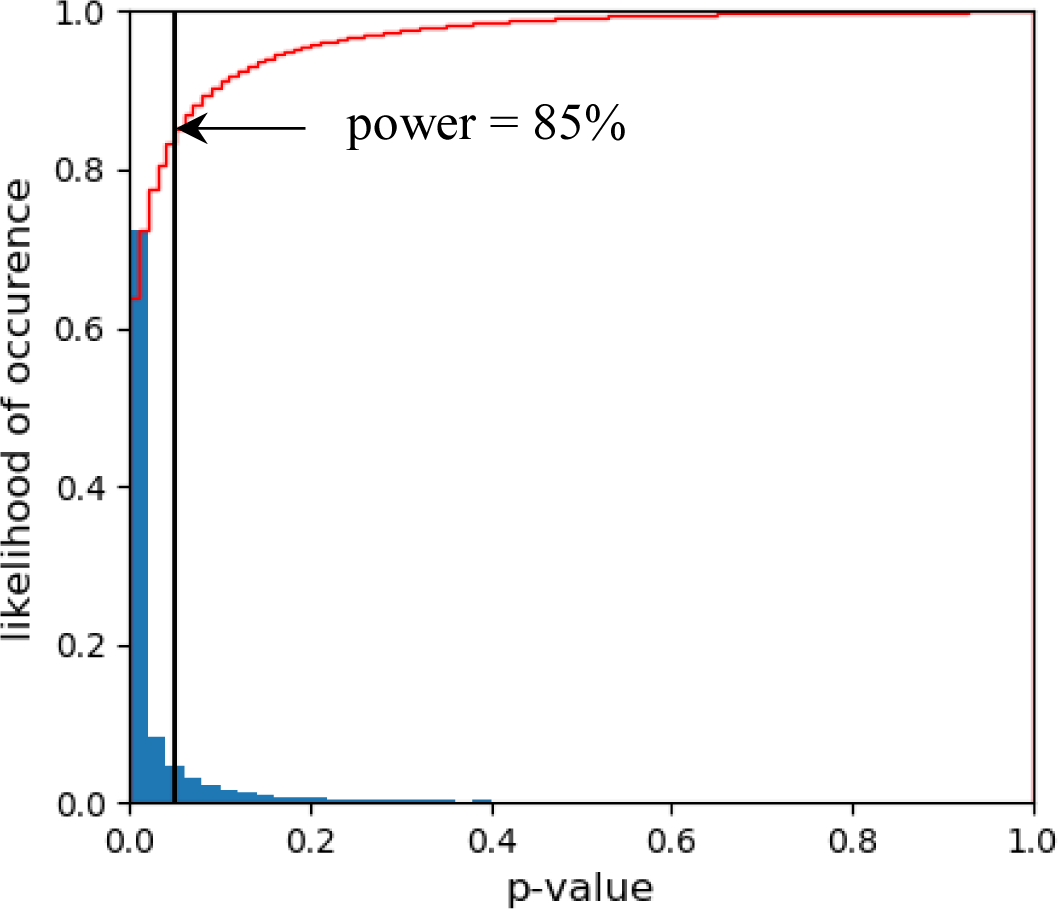
Histogram of p-values of quadratic fit on 10000 random subsamples (N=188) drawn from a simulated population (N=5227) based on the regression parameters reported by Duell et al. (2018). The blue columns represent the density of the p-values. The red line represents the cumulative histogram of the p-values. The vertical black line represents the p-value of 0.05. Note that the cumulative likelihood of p-values is above 0.8 at the intersection with the p=0.05 threshold. This implies that the probability of detecting a quadratic pattern with an alpha level of 0.05 was higher than 80%.

## References

Aklin, W. M., Lejuez, C. W., Zvolensky, M. J., Kahler, C. W., & Gwadz, M. (2005). Evaluation of behavioral measures of risk taking propensity with inner city adolescents. Behaviour research and therapy, 43(2), 215–228.

Baddeley, A. (1992). Working memory. Science, 255(5044), 556–559.

Boyer T. W. (2006). The development of risk-taking: A multi-perspective review. Developmental Review, 26(3), 291–345.

Braams, B. R., van Duijvenvoorde, A. C., Peper, J. S., & Crone, E. A. (2015). Longitudinal changes in adolescent risk-taking: a comprehensive study of neural responses to rewards, pubertal development, and risk-taking behavior. Journal of Neuroscience, 35(18), 7226–7238.

Case, R., Kurland, D. M., & Goldberg, J. (1982). Operational efficiency and the growth of short-term memory span. Journal of experimental child psychology, 33(3), 386–404.

Casey B. J., Jones R. M., Somerville L. H. (2011). Braking and accelerating of the adolescent brain. Journal of Research on Adolescence, 21, 21–33.

Chein, J., Albert, D., O’Brien, L., Uckert, K., & Steinberg, L. (2011). Peers increase adolescent risk taking by enhancing activity in the brain’s reward circuitry. Developmental Science, 14(2), 1–10.

Collado A., Felton J. W., MacPherson L.és Lejuez C. W. (2014). Longitudinal trajectories of sensation seeking, risk-taking propensity, and impulsivity across early to middle adolescence. Addictive Behaviors, 39, 1580–1588.

Collado-Rodriguez A., MacPherson L., Kurdziel G., Rosenberg L. A., & Lejuez C.W. (2014). The Relationship between Puberty and Risk-taking in the Real World and in the Laboratory. Personality and Individual Differences, 68, 143–148.

Crone E., Bullens L., Van der Plas E., Kijkuit E., Zelazo P. (2008). Developmental changes and individual differences in risk and perspective taking in adolescence. Developmental Psychopathology, 20, 1213–1229.

Defoe I. N., Dubas J. S., Figner B., van Aken M. A. G. (2015). A meta-analysis on age differences in risky decision making: adolescents versus children and adults. Psychological Bulletin, 141, 48–84.

Duell, N., Steinberg, L., Chein, J., Al-Hassan, S. M., Bacchini, D., Lei, C.,… & Lansford, J. E. (2016). Interaction of reward seeking and self-regulation in the prediction of risk taking: A cross-national test of the dual systems model. Developmental psychology, 52(10), 1593–1605.

Duell, N., Steinberg, L., Icenogle, G., Chein, J., Chaudhary, N., Di Giunta, L.,… & Pastorelli, C. (2018). Age patterns in risk taking across the world. Journal of youth and adolescence, 47(5), 1052–1072.

Eisner, M. (2002). Crime, problem drinking, and drug use: Patterns of problem behavior in cross-national perspective. The Annals of the American Academy of Political and Social Science, 580, 201–225.

Everitt, B. J., & Robbins, T. W. (2005). Neural systems of reinforcement for drug addiction: from actions to habits to compulsion. Nature neuroscience, 8(11), 1481–1489.

Fein, G., & Chang, M. (2008). Smaller feedback ERN amplitudes during the BART are associated with a greater family history density of alcohol problems in treatment-naïve alcoholics. Drug and Alcohol Dependence, 92, 141–148.

Fekete R., Filep O., Gyüre T., Ujvári K., Janacsek K. & Németh D. (2010). The examination of development of the working memory: New Hungarian standardised procedures. In: Németh D., Harsányi Sz. G. & Szokolszky Á. (Eds.). Psychological Studies –Szeged 2010) (pp. 123–132), JGYTF, Szeged

Frey, R., Pedroni, A., Mata, R., Rieskamp, J., & Hertwig, R. (2017). Risk preference shares the psychometric structure of major psychological traits. Science advances, 3(10), e1701381.

Galvan, A., Hare, T., Voss, H., Glover, G., & Casey, B. J. (2007). Risk-taking and the adolescent brain: who is at risk? Developmental science, 10(2), 8–14.

Gladwin, T. E., Figner, B., Crone, E. A., & Wiers, R. W. (2011). Addiction, adolescence, and the integration of control and motivation. Developmental Cognitive Neuroscience, 1, 364–376.

Greenhouse, S. W., & Geisser, S. (1959). On methods in the analysis of profile data. Psychometrika, 24(2), 95–112.

Griffiths, T. L., Kemp, C., & Tenenbaum, J. B. (2008). Bayesian models of cognition. In Ron Sun (ed.), Cambridge Handbook of Computational Cognitive Modeling. Cambridge University Press.

Helfinstein, S. M., Schonberg, T., Congdon, E., Karlsgodt, K. H., Mumford, J. A., Sabb, F. W.,… & Poldrack, R. A. (2014). Predicting risky choices from brain activity patterns. Proceedings of the National Academy of Sciences, 111(7), 2470–2475.

Huizenga, H. M., Crone, E. A., & Jansen, B. J. (2007). Decision-making in healthy children, adolescents and adults explained by the use of increasingly complex proportional reasoning rules. Developmental science, 10(6), 814–825.

Humphrey G. & Dumontheil I. (2016). Development of risk-taking, perspective-taking, and inhibitory control during adolescence. Developmental Neuropsychology, 41(1-2), 59–76.

Humphreys, K. L., & Lee, S. S. (2011). Risk taking and sensitivity to punishment in children with ADHD, ODD, ADHD+ ODD, and controls. Journal of Psychopathology and Behavioral Assessment, 33(3), 299–307.

Hunt, M. K., Hopko, D. R., Bare, R., Lejuez, C. W., & Robinson, E. V. (2005). Construct validity of the balloon analog risk task (BART) associations with psychopathy and impulsivity. Assessment, 12(4), 416–428.

Isaacs, E. B., & Vargha-Khadem, F. (1989). Differential course of development of spatial and verbal memory span: A normative study. British Journal of Developmental Psychology, 7(4), 377–380.

Kahneman, D. & Tversky, A. (1979). Prospect theory: An analysis of decisions under risk. Econometrica, 47, 263–291.

Kardos, Z., Kóbor, A., Takács, A., Tóth, B., Boha, R., File, B., & Molnár, M. (2016). Age-related characteristics of risky decision-making and progressive expectation formation. Behavioural Brain Research, 312, 405–414.

Kessels, R. P., Van Zandvoort, M. J., Postma, A., Kappelle, L. J., & De Haan, E. H. (2000). The Corsi block-tapping task: standardization and normative data. Applied neuropsychology, 7(4), 252–258.

Kessler, L., Hewig, J., Weichold, K., Silbereisen, R. K., & Miltner, W. H. (2017). Feedback negativity and decision-making behavior in the Balloon Analogue Risk Task (BART) in adolescents is modulated by peer presence. Psychophysiology, 54(2), 260–269.

Kóbor A., Takács Á., Janacsek K., Németh D., Honbolygó F., Csépe V. (2015). Different strategies underlying uncertain decision making: higher executive performance is associated with enhanced feedback-related negativity. Psychophysiology, 52(3), 367–377.

Koscielniak, M., Rydzewska, K., & Sedek, G. (2016). Effects of age and initial risk perception on balloon analog risk task: The mediating role of processing speed and need for cognitive closure. Frontiers in Psychology, 7(659).

Lange-Küttner, C., Averbeck, B. B., Hirsch, S. V., Wießner, I., & Lamba, N. (2012). Sequence learning under uncertainty in children: self-reflection vs. self-assertion. Frontiers in psychology, 3, 127.

Lee, M. D., & Wagenmakers, E.-J. (2013). Bayesian cognitive modeling: A practical course. Cambridge University Press.

Lejuez, C. W., Read, J. P., Kahler, C. W., Richards, J. B., Ramsey, S. E., Stuart, G. L., & Brown, R. A. (2002). Evaluation of a behavioral measure of risk-taking: The Balloon Analogue Risk Task (BART). Journal of Experimental Psychology: Applied, 8, 75–84.

Lejuez, C. W., Aklin, W. M., Zvolensky, M. J., & Pedulla, C. M. (2003). Evaluation of the Balloon Analogue Risk Task (BART) as a predictor of adolescent real-world risk-taking behaviours. Journal of Adolescence, 26(4), 475–479.

MacPherson L, Magidson J, Reynolds EK, Kahler CW, Lejuez CW. (2010) Changes in sensation seeking and risk-taking propensity predict increases in alcohol use among early adolescents. Alcoholism: Clinical and Experimental Research, 34, 1400–1408.

McCormick, E. M., Qu, Y., & Telzer, E. H. (2017). Activation in context: differential conclusions drawn from cross-sectional and longitudinal analyses of adolescents’ cognitive control-related neural activity. Frontiers in human neuroscience, 11, 141.

Paulsen D., Platt M., Huettel S.A., & Brannon E.M. (2011). Decision-making under risk in children, adolescents, and young adults. Frontiers in Psychology, 2(72).

Peper, J. S., Braams, B. R., Blankenstein, N. E., Bos, M. G., & Crone, E. A. (2018). Development of Multifaceted Risk Taking and the Relations to Sex Steroid Hormones: A Longitudinal Study. Child development, 89(5), 1887–1907.

Racsmány, M., Lukács, Á., Németh, D., & Pléh, C. (2005). Hungarian diagnostic tools of verbal working memory functions. Magyar Pszichológiai Szemle (Hungarian Psychological Review), 4(60), 479–506.

Reyna, V.F., Farley, F. (2006). Risk and rationality in adolescent decision making –implications for theory, practice, and public policy. Psychological science in the public interest, 7(1), 1–44.

Reyna, V. F., & Rivers, S. E. (2008). Current theories of risk and rational decision making. Developmental Review, 28(1), 1–11.

Schmitz, F., Manske, K., Preckel, F., & Wilhelm, O. (2016). The multiple faces of risk-taking. European Journal of Psychological Assessment, 32(1), 17–38.

Shulman, E. P., & Cauffman, E. (2014). Deciding in the dark: Age differences in intuitive risk judgment. Developmental psychology, 50(1), 167–177.

Smith, A. R., Chein, J., & Steinberg, L. (2014). Peers increase adolescent risk taking even when the probabilities of negative outcomes are known. Developmental Psychology, 50(5), 1564–1568.

Somerville, L. H., & Casey, B. J. (2010). Developmental neurobiology of cognitive control and motivational systems. Current Opinion in Neurobiology, 20, 236–241.

Spreen, O., & Strauss, E. (1991). Controlled oral word association (word fluency).

Spreen O, Strauss E. A compendium of neuropsychological tests. Oxford: Oxford University Press, 219–27.

Steinberg, L. (2004). Risk-taking in adolescence: what changes, and why? Annals of the New York Academy of Sciences, 1021, 51–58.

Steinberg L. (2007). Risk-taking in adolescence: New perspectives from brain and behavioral science. Current Directions in Psychological Science, 16, 55–59.

Steinberg L., Albert D., Cauffman E., Banich M., Graham S., & Woolard J. (2008). Age differences in sensation seeking and impulsivity as indexed by behavior and self-report: evidence for a dual systems model. Developmental Psychology, 44(6), 1764–1778.

Steinberg L. (2010). A dual systems model of adolescent risk-taking. Developmental Psychobiology, 52, 216–224.

Sweeten, G., Piquero, A. R., & Steinberg, L. (2013). Age and the explanation of crime, revisited. Journal of youth and adolescence, 42(6), 921–938.

Tanczos, T., Janacsek, K., Nemeth, D. (2014a). Verbal fluency tasks I. Investigation of the Hungarian version of the letter fluency task between 5 and 89 years of age. Psychiatria Hungarica, 29(2), 158–180.

Tanczos, T., Janacsek, K., Nemeth, D. (2014b) Verbal fluency tasks II. Investigation of the Hungarian version of the semantic fluency task between 5 and 89 years of age. Psychiatria Hungarica, 29(2), 181–207.

Takács Á., Kóbor A., Janacsek K., Honbolygó F., Csépe V., Németh D. (2015). High trait anxiety is associated with attenuated feedback-related negativity in risky decision making. Neuroscience Letters, 600, 188–192.

Tukey, J. W. (1977). Exploratory Data Analysis. Addison-Wesley.

Van Duijvenvoorde, A. C., Jansen, B. R., Bredman, J. C., & Huizenga, H. M. (2012). Age-related changes in decision making: Comparing informed and noninformed situations. Developmental Psychology, 48(1), 192–203.

Van Leijenhorst, L., Westenberg, P. M., & Crone, E. A. (2008). A developmental study of risky decisions on the cake gambling task: age and gender analyses of probability estimation and reward evaluation. Developmental Neuropsychology, 33(2), 179–196.

Van Leijenhorst, L., Moor, B. G., de Macks, Z. A. O., Rombouts, S. A., Westenberg, P. M., & Crone, E. A. (2010). Adolescent risky decision-making: neurocognitive development of reward and control regions. Neuroimage, 51(1), 345–355.

Weller J., Levin I., Denburg N. (2010). Trajectory of risky decision making for potential gains and losses from ages 5 to 85. J. Behavioral Decision Making, 24(4), 331–344.

Willoughby, T., Good, M., Adachi, P. J., Hamza, C., & Tavernier, R. (2014). Examining the link between adolescent brain development and risk taking from a social–developmental perspective (reprinted). Brain and cognition, 89, 70–78.

